# A functional amyloid scaffold shapes insect egg coats

**DOI:** 10.64898/2026.05.29.727653

**Authors:** Katerina Konstantoulea, Peter Kunach, Harichandra Tagad, Marc I. Diamond, Nikolaos N. Louros

## Abstract

Functional amyloids serve as structural scaffolds across biology, yet the molecular architecture and assembly principles of many remain unresolved. The lepidopteran egg coat, or chorion, presents a striking example: hundreds of paralogous proteins sharing a conserved central domain form a mechanically resilient amyloid matrix essential for embryo protection. Here, combining evolutionary analysis of more than 500 sequences with cryo-electron microscopy and biophysical assays, we determine the atomic structure of chorion amyloid filaments and uncover the principles governing their assembly. Contrary to previous structural predictions, chorion filaments adopt a β-serpentine fold stabilized by a short hexapeptide motif that forms homotypic steric-zipper interfaces, self-assembles autonomously, and seeds full-length filament growth. Amyloid formation proceeds through secondary nucleation, while thermodynamic and structural analyses support a hierarchical assembly mechanism in which motif-driven interactions nucleate filament formation prior to consolidation of the mature yet lalbile protofilament core. These findings establish the molecular basis of insect egg-coat assembly and demonstrate that mechanisms commonly associated with pathological aggregation can also operate within a biologically regulated functional amyloid framework.

## Introduction

Amyloid filaments, defined by their cross-β architecture^1^, have emerged as a versatile structural solution that combines high mechanical stability with resistance to chemical and proteolytic degradation^2-6^, and are deployed across kingdoms of life to build mechanically robust extracellular biomaterials^2,3,6^. Despite being primarily associated with neurodegenerative and other pathological conditions^4,7^, an expanding number of amyloids have been identified to catalyse various biological functions, including pigmentation^8-10^, signalling^11-15^, hormone storage^16,17^, extracellular scaffolding^18-22^, and antimicrobial defence^23-28^. A key distinction between functional and pathological amyloids lies in their regulation: functional amyloid formation is typically subject to tight spatiotemporal control^2,29^, and the resulting assemblies are, in certain cases, reversible^30^. Their assembly mechanisms have also been assumed to differ, as functional amyloids are generally reported to assemble through primary nucleation or elongation of their native fold, whereas secondary nucleation, a surface-catalysed process that amplifies aggregation in cases of pathological amyloids, has rarely been described in functional contexts^19^.

Egg coats represent a striking example of functional amyloid-based extracellular scaffolds. In vertebrates, oocytes are enclosed by protective envelopes as filamentous extracellular matrices with amyloid properties that are essential for oocyte protection and fertilization^31-35^. Insects have converged on an analogous strategy. The lepidopteran egg coat, or chorion, constitutes the outermost protective layer of the eggshell and withstands substantial mechanical stress, desiccation, and environmental exposure while remaining permeable to gas exchange^36^. Chorion assembly occurs late in oogenesis and involves hundreds of homologous proteins that organize into a highly ordered, insoluble matrix with a lamellar ultrastructure of packed amyloid filaments^37,38^. Biochemical, spectroscopic, and ultrastructural studies have established the amyloid features of the chorion^38-42^, yet the molecular architecture and assembly principles underlying chorion filament formation have remained unknown.

A defining feature of lepidopteran chorion proteins is a conserved glycine-rich central domain shared across two homologous protein families^43,44^. An archetype protein sequence representing the central domain of these families (cA) readily assembles into amyloid filaments, establishing the intrinsic amyloidogenic properties of the central domain^38,45^. Prior modelling studies suggested that this domain might adopt a β-solenoid-like fold^45^, a compact architecture seen in several other functional amyloids^46-51^. Yet chorion proteins are encoded by hundreds of paralogous genes within and across lepidopteran species^43^, raising a fundamental question: how does such a diverse protein ensemble encode a common supramolecular architecture capable of forming a coherent extracellular matrix?

Here, we integrate evolutionary analysis, biophysical characterisation, and cryo-electron microscopy (cryo-EM) to define the molecular architecture of insect chorion amyloids. By analysing more than 500 chorion sequences across Lepidoptera, we show that the conserved central domain occupies a constrained region of sequence space and that the cA archetype captures its defining sequence features. Unexpectedly, cryo-EM reveals that cA filaments adopt a β-serpentine fold stabilised by recurrent steric-zipper interfaces, including a homotypic zipper formed by a short hexapeptide motif. This motif self-assembles autonomously, adopts a steric-zipper geometry matching that observed within the full-length filaments, and promotes cross-seeding of the chorion archetype. Assembly is dominated by secondary nucleation, and thermodynamic profiling supports a hierarchical model in which motif-driven nucleation precedes consolidation of the mature protofilament core. Together, these findings provide insight into the molecular architecture of insect egg coat amyloids and reveal how assembly mechanisms more commonly associated with pathological aggregation can operate within an evolutionarily constrained and biologically regulated functional context.

## Results

### Evolutionary analysis reveals deep sequence conservation of chorion proteins

To characterize the evolutionary logic underpinning the Lepidopteran chorion family, we assembled and aligned the conserved central domains of proteins across 32 species spanning the major Lepidopteran lineages (**Fig. 1A**). Each chorion protein contains a single copy of this central domain with a conserved length of 51 amino acids, while flanking regions vary extensively in both length and sequence composition between family members. Phylogenetic analysis revealed species-resolved groupings, with expansions of homologous chorion repertoires in *B. mori* and *M. sexta*, alongside more diffuse branching among butterfly homologs. A species-wise protein count distribution highlighted this quantitatively, with *B. mori* contributing the largest repertoire, whereas other species were represented by smaller sequence sets (**Fig. 1B**). Multiple sequence alignment of the central domain revealed strong positional conservation across its length, with a highly regular glycine-rich pattern interspersed primarily with short hydrophobic residues, as reported by earlier studies^52^. Despite the extensive paralogs across species, no internal insertions or deletions were observed within the conserved central domain. On the basis of this alignment, we derived a consensus sequence capturing the residue frequency at each position, matching the previously described cA archetype shown to form amyloid filaments^38,45^.

**Figure 1.**
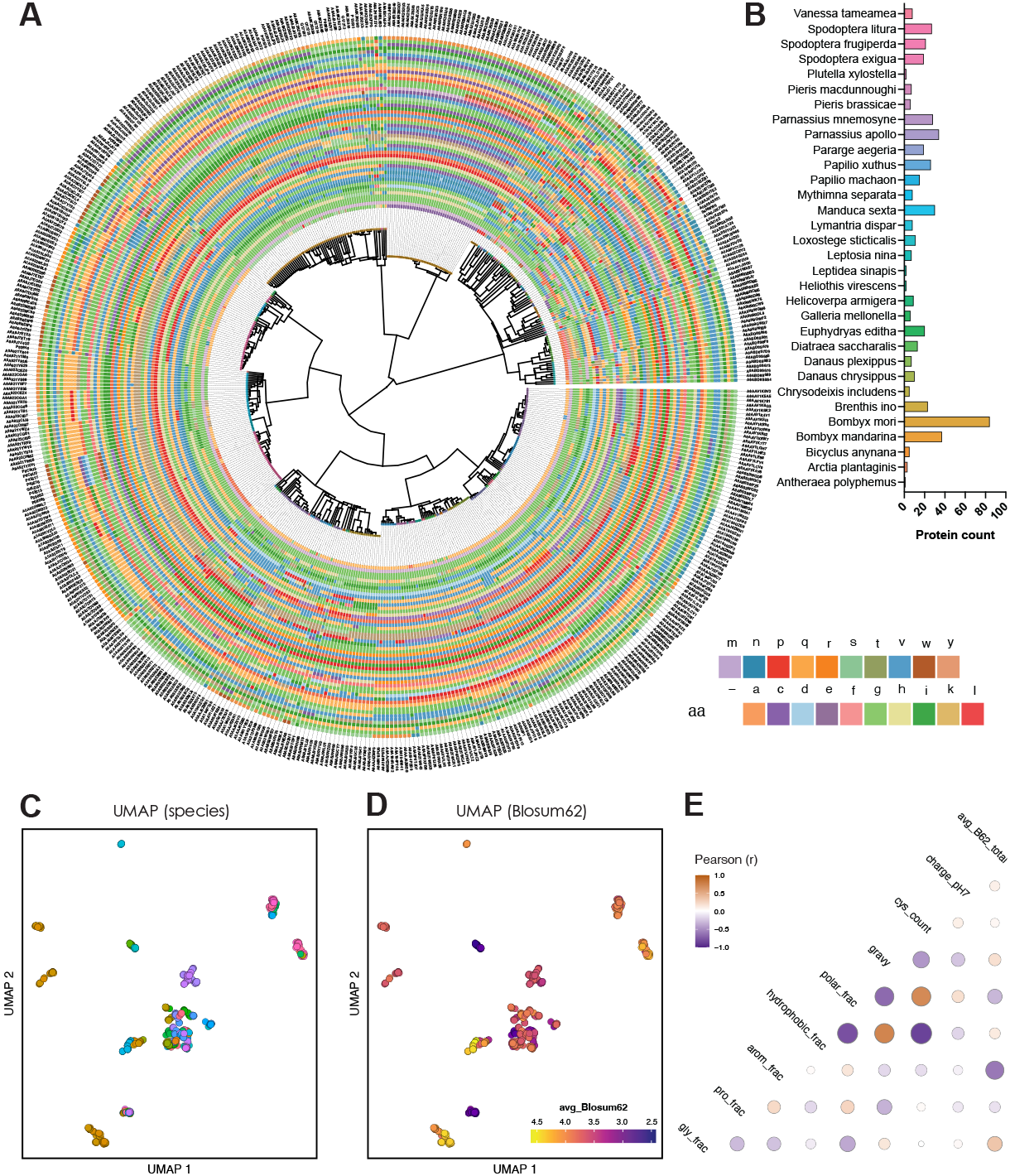
Evolutionary compression of Lepidopteran chorion proteins. (**A**) Circular maximum-likelihood phylogeny of the conserved central domains from chorion proteins across Lepidopteran species. **(B)** Distribution of protein counts per species used in the evolutionary analysis. **(C)** UMAP projection of pairwise sequence dissimilarities coloured by organism. **(D)** UMAP coloured by average BLOSUM62 similarity to the archetype, showing strong sequence convergence. **(E)** Correlation matrix of sequence-derived properties versus average BLOSUM62 similarity.

To determine how these sequences populate sequence space, we computed pairwise sequence dissimilarities and embedded them using uniform manifold approximation and projection (UMAP). Colouring the embeddings by species revealed a sequence space with a central cluster and several satellites, including multiple *B. mori*-enriched groups and more distant butterfly-dominated regions (**Fig. 1C**). Colouring the same projection by average BLOSUM62 similarity to the archetype, a metric that captures evolutionary distance, revealed strong convergence across sequences (**Fig. 1D**). Most sequences exhibited high similarity scores (avg_Blosum62 > 3.5), while the distal satellite clusters retained substantial similarity (avg_Blosum62 > 2.5), indicating that even the most divergent sequences remain close to the archetype. Correlation analysis of sequence-derived physicochemical features revealed linked compositional constraints (**Fig. 1E**): glycine fraction, hydropathy (GRAVY), and fraction of hydrophobic residues covary strongly with archetype similarity, while aromatic content and fraction of polar residues are anti-correlated. These trends suggest that the archetype is a representative baseline of this protein family and indicate selective pressure to preserve flexibility and steric compatibility within the conserved domain while disfavouring bulky side chains.

### Assembly mechanism and stability of chorion amyloid filaments

To characterise the assembly behaviour of the chorion archetype, we first confirmed that it assembles into amyloid filaments. Transmission electron microscopy (TEM) of end-point reactions confirmed the formation of twisting amyloid filaments, consistent with canonical cross-β assemblies (**Fig. 2A**). To quantify filament formation kinetics, we monitored assembly using thioflavin-T (ThT) fluorescence across a range of monomer concentrations (4 - 110 µM) (**Fig. 2B-C**). Global fitting of the kinetic traces was performed to assess the relative contributions of primary (**Fig. 2B**) and secondary nucleation (**Fig. 2C**) in filament growth. A model dominated by secondary nucleation provided the best fit to the experimental data, indicating that filament growth is driven by surface-catalysed generation of new aggregates rather than by a nucleation-elongation model. Plotting the reaction half-times (t_1/2_) against the initial monomer concentrations further supports this. The linear fit of the double logarithmic plot indicates the absence of saturating microscopic steps involved over the specific concentration range (**Fig. 2D**), while the calculated scaling exponent (γ ∼ -0.44) matches reported slopes of faster secondary nucleation-driven aggregation^53,54^. Such kinetic behaviour has been associated with pathogenic amyloids^19^.

**Figure 2.**
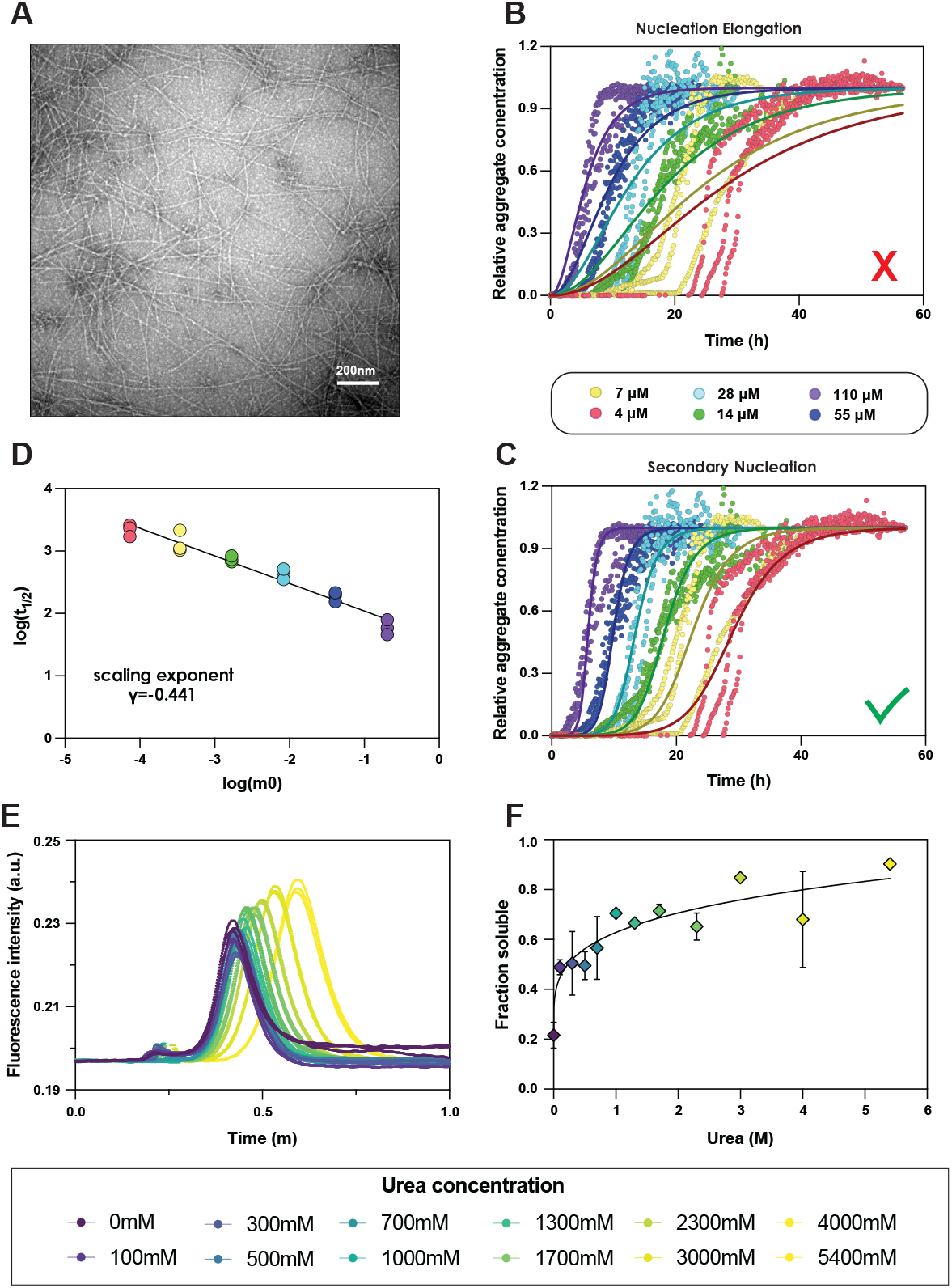
Properties of amyloid filaments formed by cA self-assembly. (**A**) Negative-stain micrographs of end-point filaments showing long, unbranched filaments with twisted morphologies, characteristic of cross-β amyloid assemblies. Scale bar, 200 nm. **(B-C)** Thioflavin-T fluorescence kinetics (n=3 independent repeats) recorded at increasing initial monomer concentrations (4-110 μM). Solid lines represent global fits to a kinetic model dominated by primary **(B)** or secondary nucleation **(C). (D)** Log-plot of aggregation half-time (t_1/2_) versus initial monomer concentration (m_0_). The linear dependence yields a scaling exponent γ = −0.441, consistent with aggregation dominated by secondary nucleation rather than primary nucleation or fragmentation. **(E)** FIDA traces of filament aliquots exposed to increasing urea concentrations (0-5.4 M). Progressive peak broadening and shifts reflect increasing solubilization of fibrillar material (n=3 independent repeats). **(F)** Soluble fraction as a function of urea concentration quantified from FIDA traces in **(D)**. Solid line represents a global fit yielding a low free energy of stabilization (ΔG ∼ -4.85 kcal mol^-1^), consistent with a thermodynamically labile functional amyloid.

To evaluate the thermodynamic stability of the assembled filaments, we performed chemical denaturation experiments monitored by fluorescence-induced dispersion analysis (FIDA). This method distinguishes soluble and fibrillar species based on their distinct transport properties in laminar flow^55,56^. FIDA analysis produced well-defined elution profiles, and quantification of the relative fractions revealed concentration-dependent chemical denaturation (**Fig. 2E**). A global nonlinear fit of all data points indicated that chorion filaments dissociate at low denaturant concentrations, yielding a ΔG value of −4.85 kcal·mol^−1^, consistent with labile amyloid assembly (**Fig. 2F**) and matching previously reported stabilities for other functional amyloid filaments^57^.

### Cryo-EM determination of chorion amyloid filaments

To define the molecular architecture of amyloid filaments formed by the chorion archetype, we determined their structures using cryo-electron microscopy. Reference-free two-dimensional (2D) classification and hierarchical clustering (**Fig. S1**) identified six fibril polymorphs, two of which converged to the same final reconstruction and were thus merged into the final Type 1 class (**Fig. 3A-B**). Type 1 represented the dominant population (∼44% of classified particles), followed by Type 3 (∼26%), Type 4 (∼15%), Type 5 (∼11%), and Type 6 (∼4%) (**Fig. 3A**). The different polymorphs displayed distinct crossover distances and filament widths, with Type 6 fibrils exhibiting very wide crossovers (∼1237 Å), Type 1, Type 4, and Type 5 fibrils exhibiting similar wide spacings (∼475 Å, ∼520 Å, and ∼584 Å, respectively), and Type 3 fibrils adopting a tighter helical twist with a crossover distance of ∼303 Å (**Fig. 3B-C**).

**Figure 3.**
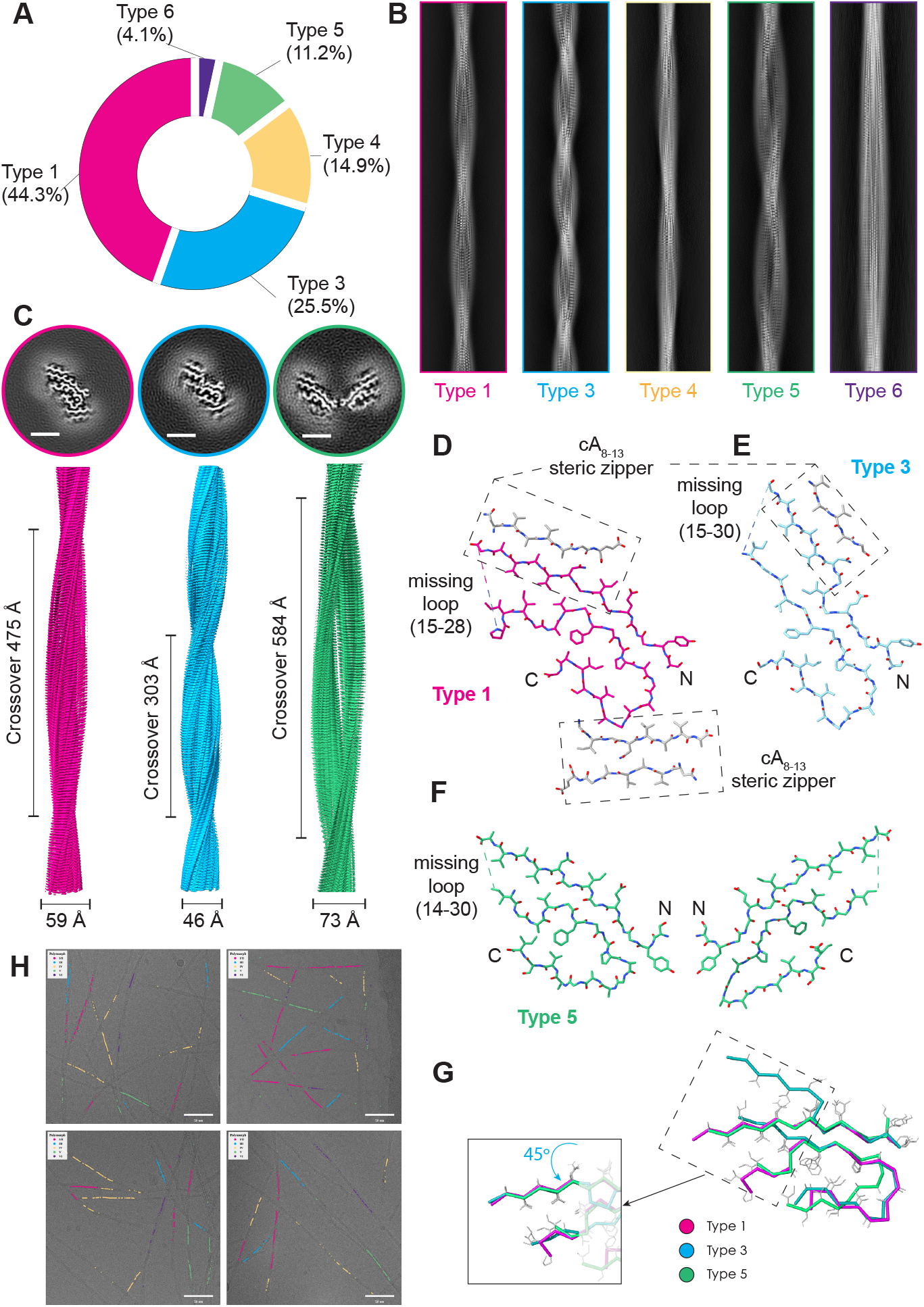
Cryo-EM structures of the chorion amyloid filaments. (**A**) Relative particle populations of the identified fibril polymorphs obtained from reference-free 2D classification. **(B)** Stitched reference-free 2D class averages corresponding to the different polymorphs. **(C)** Representative cryo-EM cross-sections and density maps of the major polymorphs showing differences in filament width and protofilament organization. Scale bar: 25Å **(D-F)** Corresponding atomic models of the Type 1 **(D)**, Type 3 **(E)**, and Type 5 **(F)** fibrils. All resolved polymorphs adopt a conserved cross-β β-serpentine architecture. cA_8-13_ motifs form homotypic steric-zipper interfaces mediating lateral contacts between adjacent filament surfaces, whereas residues 15–28 remain unresolved in all structures. **(G)** Superposition of the Type 1, Type 3, and Type 5 atomic models reveals extensive conservation of the β-serpentine backbone scaffold across polymorphs, with the 8-13 and 30-33 segments forming a shared steric zipper interfaces that is angled by 45° in Type 3 filaments. **(H)** Mapping of classified particle segments onto representative cryo-EM micrographs reveals coexistence of multiple polymorphs within the same fibrillar assemblies.

Three-dimensional reconstruction yielded high-resolution density maps and atomic models for Type 1, Type 3, and Type 5 filaments (**Fig. 3D-F**), while detailed atomic interpretation was precluded in Type 4 and Type 6. Despite differences in filament morphology and helical symmetry, the resolved polymorphs converged onto a conserved cross-β architecture with a β-serpentine backbone trajectory. In all conformers, residues 15-28 formed a disordered loop that was absent from the ordered fibril core (**Fig. 3D-F**). Superposition of the Type 1, Type 3, and Type 5 atomic models revealed extensive structural conservation across the ordered regions of the fibrils (**Fig. 3G**). The overall backbone trajectory was overlapping between conformers, indicating that the polymorphs preserve a common structural scaffold rather than representing fundamentally distinct fibril folds. Structural differences were localised primarily to glycine-centred backbone reorientations while maintaining the global β-serpentine architecture. Notably, residues 1-6 formed a conserved heterotypic steric zipper with residues 35-40 across all resolved structures, whereas residues 8-13 pack against residues 30-33. The recurrent involvement of the former motif (cA_8-13_) was a particularly prominent feature of the fibril structures. In both Type 1 and Type 3 filaments, this motif formed additional face-to-face homotypic steric zipper interfaces mediating lateral interactions with the main protofilament (**Fig. 3D-E, shown in grey**). Additional peripheral densities compatible with similar cA_8-13_-mediated interactions were also observed in Type 5 filaments, although these regions were less well resolved, suggesting that cA_8-13_-mediated interactions represent a conserved feature of chorion fibril assembly.

To investigate the spatial organization of the different polymorphs, classified particle segments were mapped back onto representative cryo-EM micrographs (**Fig. 3H**). Distinct polymorphs frequently coexisted within the same fibrillar networks and, in multiple cases, were observed along apparently continuous filament trajectories. Together with the extensive structural similarity revealed by backbone superposition, these observations suggest that the identified polymorphs do not represent fully independent strain-like architectures but rather closely related states within a shared conformational landscape. The conserved β-serpentine scaffold supports a model in which chorion filaments sample structurally adjacent packing states while preserving a common underlying fold.

### A hexapeptide motif encodes an autonomous amyloid nucleus

The repeated appearance of the cA_8-13_ steric zipper interface suggested it may represent an intrinsic self-assembling module. We therefore synthesized (**Fig. S2**) and tested whether this motif recapitulates the chorion fibril zipper geometry in isolation and whether it can seed the assembly of the archetype into filaments. Electron micrographs revealed the formation of abundant unbranched amyloid filaments of the isolated peptide (**Fig. 4A**). In contrast to cA filaments, these peptide assemblies were ThT-negative (**Fig. S3**). To circumvent this, we assessed amyloid formation using a luminescent oligothiophene probe (pFTAA), which readily bound the peptide filaments, supporting the formation of ordered β-rich aggregates (**Fig. 4B**). Because pFTAA acts as a conformation-sensitive reporter of amyloid architecture and has been widely used to probe fibril heterogeneity, strain-specific structures, and fibril remodeling^58-64^, we next performed spectral analysis of emission profiles followed by principal component analysis (PCA). This revealed superimposable spectra obtained from cA_8-13_ peptide filaments and archetype filaments (**Fig. 4C**), suggesting that pFTAA engages related supramolecular environments in the two assemblies, consistent with shared structural features between filaments. To further contextualize these observations, we extended the analysis to other canonical amyloid fibrils, including Aβ_42_, tau, TAF15, and α-synuclein. PCA of the corresponding pFTAA emission spectra demonstrated that archetype and cA_8-13_ filaments cluster together in eigenspace, while clearly separating from all other amyloid species (**Fig. 4C**). This segregation indicates that pFTAA reports on a shared binding environment specific to the chorion-derived assemblies, further supporting structural similarity between the peptide and archetype filaments, and distinguishing them from other amyloid architectures.

**Figure 4.**
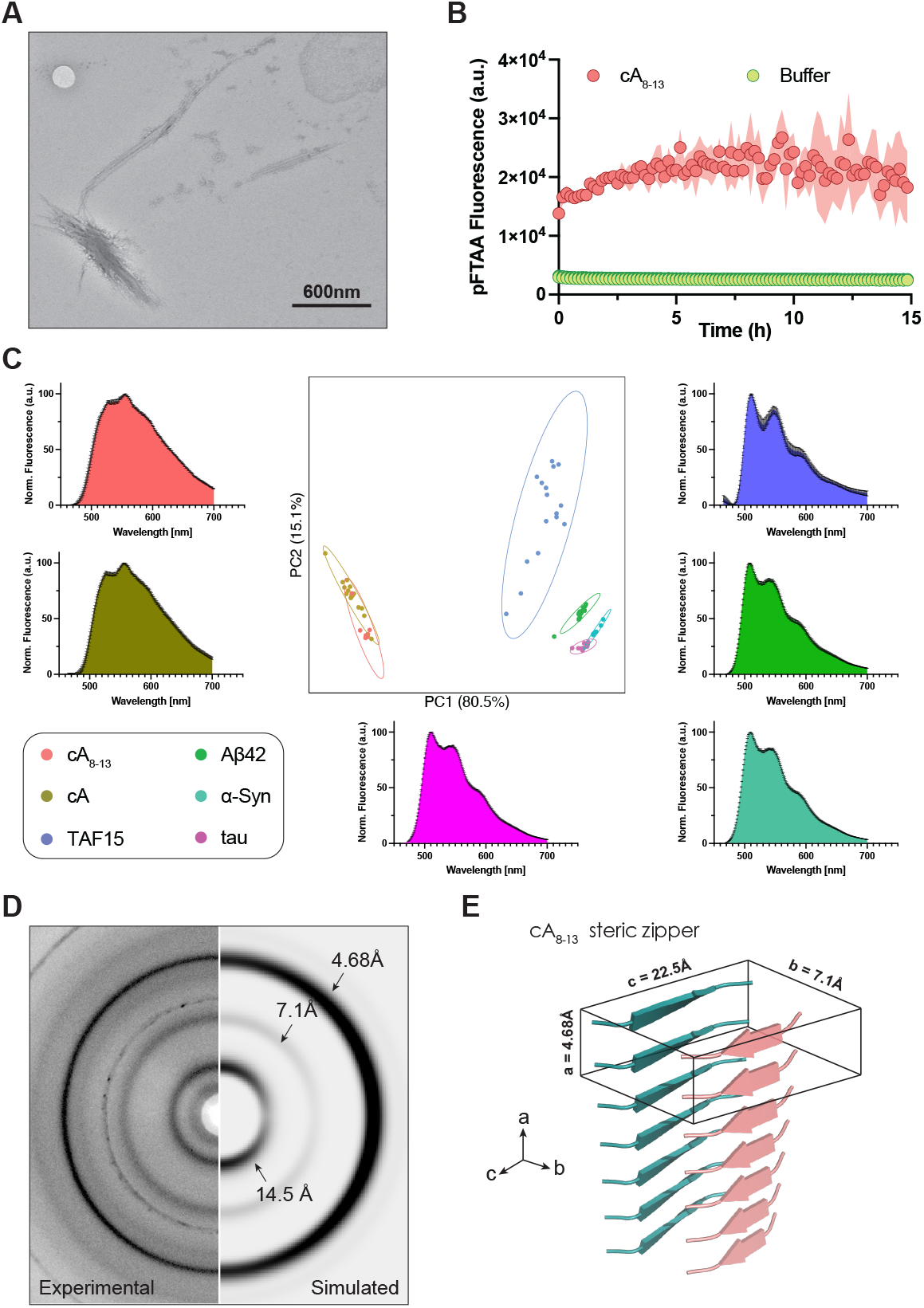
Amyloid filaments formed by the cA_8-13_ amyloid motif. (**A**) Negative-stain micrograph of filaments formed by the isolated hexapeptide, showing unbranched amyloid-like assemblies. Scale bar, 600 nm. **(B)** Fluorescence aggregation kinetics of cA_8-13_. Shaded regions represent standard deviation (n=3 independent repeats). **(C)** Principal component analysis of pFTAA fluorescence emission spectra collected from filaments formed by cA_8-13_ and the cA archetype, alongside representative pathological amyloids (Aβ42, α-synuclein, tau, and TAF15). Data are presented as mean ± SD from independent experiments (cA, n = 16; cA_8-13_, n = 7; Aβ_42_, n = 16; tau, n = 13; TAF15, n = 16; α-synuclein, n = 6). **(D)** ray fibre diffraction patterns of cA_8-13_ filaments compared with simulated diffraction derived from the steric-zipper interface extracted from the Type 1 filaments. Characteristic reflections at ∼4.7 Å, ∼7.1 Å, and ∼14.5 Å are indicated. **(E)** Atomic model of the cA_8-13_ steric zipper and the corresponding ortho-rhombic unit cell used for diffraction simulations.

To further assess the structural relationship between the motif filaments and the steric-zipper interfaces observed in the cryo-EM structures, we performed X-ray fibre diffraction on the isolated peptide assemblies. The resulting diffraction patterns showed the characteristic ∼4.7 Å meridional reflection consistent with β-strand spacing, alongside additional equatorial reflections at ∼7.1 Å and ∼14.5 Å (**Fig. 4D**), with the former spacing matching the inter-β-strand distances observed within the homotypic steric-zipper interfaces formed by the cA_8-13_ motif in the cryo-EM structures (**Fig. 4E**). Using the atomic coordinates of this isolated steric-zipper interface, extracted from the Type 1 fibril model, we simulated diffraction patterns that closely reproduced the experimental patterns when modelled using an orthorhombic unit cell accommodating stacked cA8-13 motifs (**Fig. 4D-E**). These results further support that the hexapeptide adopts in isolation a steric-zipper packing geometry similar to that formed within the chorion filaments.

We next asked whether this autonomous steric-zipper module can influence the assembly of the archetype. Addition of preformed cA_8-13_-derived filaments to monomeric protein shortened the lag phase of aggregation, but with lower efficiency compared to self-seeding (**Fig. 5A-C**). These results indicate that the cA_8-13_ zipper can further act as an early structural template that promotes seeded propagation of the chorion archetype sequence.

**Figure 5.**
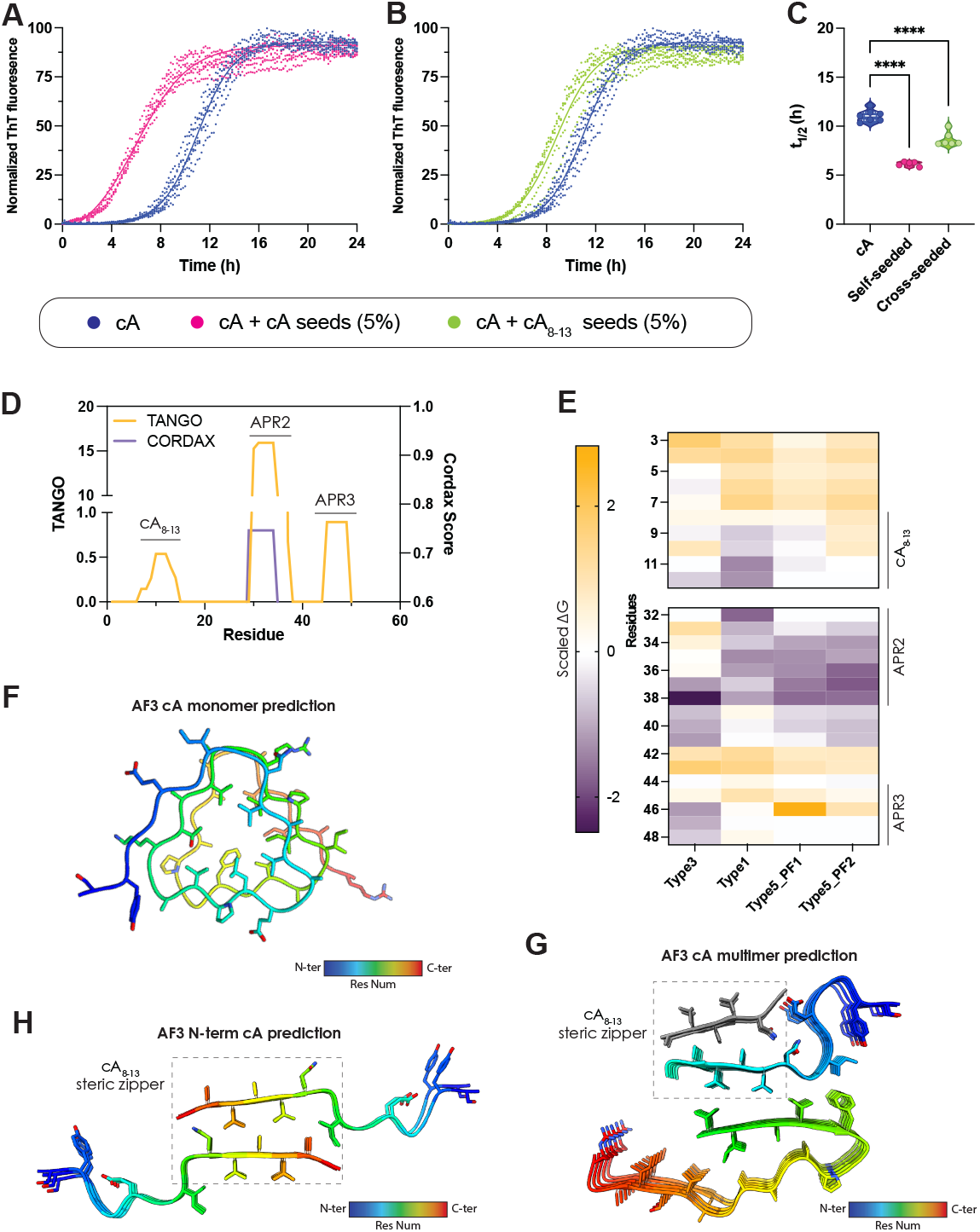
Impact of cA_8-13_ motif on cA amyloid assembly. (**A**) Fluorescence aggregation kinetics of cA in the absence and presence of homologous cA seeds (5% w/w), showing accelerated assembly upon self-seeding. **(B)** Fluorescence aggregation kinetics of cA in the absence and presence of preformed cA_8-13_ filaments (5% w/w). **(C)** Aggregation half-times extracted from (A-B). Statistics: One-way ANOVA with multiple t-test comparison. **(D)** Sequence-based aggregation propensity predictions identifying the N-terminal cA_8-13_ segment and two additional aggregation-prone regions (APR2, APR3). **(E)** Thermodynamic profiling across polymorphs displayed as a residue-by-polymorph heatmap, highlighting systematic shifts in stabilization. **(F)** AF3 prediction for monomeric cA, yielding a compact β-solenoid fold prediction. **(G)** AF3 multimer prediction for cA, producing an amyloid-like assembly, with additional cA_8-13_ homotypic steric-zipper contacts (highlighted in grey). **(H)** AF3 multimer prediction of an N-terminal cA fragment (1-14), prioritizing a homotypic cA_8-13_ steric zipper, consistent with an early self-complementary interaction. AF models are coloured-coded from N-to C-terminus.

### Modular interfaces shape chorion filament assembly

Sequence-based aggregation predictors identified cA_8-13_ as a major aggregation-prone region (APR), together with two additional segments spanning residues 32–38 and 45–50 (**Fig. 5D**). In the cryo-EM structures, the latter two segments form a heterotypic steric-zipper interface within the central protofilament core, suggesting that additional stabilizing interactions emerge in the context of the extended β-serpentine scaffold. Consistent with this modular organization, thermodynamic profiling of the resolved polymorphs revealed systematic energetic signatures between Type 1, Type 3, and Type 5 filaments (**Fig. 5E**). Type 3 filaments displayed strong localized energetic hotspots, whereas in Type 1 and Type 5 stabilization became progressively redistributed across the central protofilament core. These observations suggest that while cA_8-13_-mediated interactions contribute strongly to filament assembly, additional aggregation-prone regions may increasingly consolidate the main protofilament architecture.

To investigate whether the lateral cA_8-13_ interfaces observed experimentally could emerge as preferred early contacts during assembly, we performed AlphaFold (AF) folding predictions using (i) a single copy of the monomeric sequence, (ii) multimeric N-terminal fragments of the archetype, and (iii) multimeric archetype assemblies supplemented with additional copies of cA_8-13_ motifs. Whereas the monomer was predicted to adopt a β-solenoid fold (**Fig. 5F**), consistent with earlier sequence-based hypotheses45, multimeric predictions converged onto fibril-like arrangements resembling, to an impressive degree, the experimentally observed βserpentine topology (**Fig. 5G**). Notably, both full-length and isolated N-terminal fragment multimeric predictions repeatedly prioritized cA8-13-mediated homotypic steric-zipper contacts (**Fig. 5H**), supporting the idea that this motif encodes a low-barrier selfcomplementary interaction that is readily sampled under oligomeric constraints.

Notably, both full-length and isolated N-terminal fragment multimeric predictions repeatedly prioritized cA_8-13_-mediated homotypic steric-zipper contacts (**Fig. 5H**), supporting the idea that this motif encodes a low-barrier self-complementary interaction that is readily sampled under oligomeric constraints.

Together with the ability of isolated cA_8-13_ assemblies to nucleate chorion filament formation, these observations support a model in which cA_8-13_ acts as an autonomous assembly module that contributes to the early organization of the fibril architecture. In this framework, the different chorion polymorphs are best interpreted not as distinct strain-like states, but as closely related packing solutions within a shared assembly landscape. The coexistence of multiple polymorphs, combined with their extensive structural conservation, suggests that chorion filaments retain substantial conformational plasticity while preserving a common β-serpentine scaffold. An alternative, non-exclusive possibility is that some polymorphs may represent partially consolidated structural states sampled during filament maturation, mirroring recent findings from complementary temporal structural studies of different amyloid proteins^65-67^. In such a scenario, the shared structural scaffold and progressive redistribution of stabilizing interactions across Type 3, Type 1, and Type 5 filaments would be consistent with increasingly consolidated protofilament architectures.

## Discussion

In this work, we set out to understand how lepidopteran chorion proteins assemble into a functional amyloid matrix and whether their assembly principles resemble those of other functional amyloids. Using an evolutionary archetype representing the conserved central domain of the chorion protein families, we show that it readily forms amyloid filaments whose kinetics are dominated by secondary nucleation. Functional amyloids are more often reported to assemble through primary nucleation, elongation, and fragmentation^19^. Instead, secondary nucleation is primarily associated with pathological amyloids such as Aβ^68^, α-synuclein^69^, tau^70^, and IAPP^71^. One explanation proposed for this distinction is architectural. Several functional amyloids adopt β-solenoid-like folds composed of imperfect repeats stacked along the fibril axis^51,72,73^. Such architectures may limit the formation of well-defined fibril surface features capable of catalysing secondary nucleation. In line with this, we previously suggested that chorion proteins might assemble through a β-solenoid stacking mechanism^45^, with monomeric AF predictions being consistent with this hypothesis (**Fig. 5F**). However, our cryo-EM structures reveal that chorion filaments instead adopt a β-serpentine architecture composed of individual monomer rungs stacked along the cross-β fibril axis. Multimeric AF predictions similarly converged to a cross-β topology, indicating that oligomeric constraints are sufficient to redirect away from a solenoid-like conformation. Strikingly, inclusion of additional copies of the cA_8-13_ hexapeptide motif in multimeric predictions strongly converges toward the β-serpentine architecture resolved by cryo-EM (**Fig. 5G**), suggesting that secondary contacts mediated by this aggregation-prone segment likely contribute to defining the final topology of chorion fibrils.

This notion is reinforced by the intrinsic assembly properties of the cA_8-13_ motif. The isolated hexapeptide readily forms amyloid filaments, consistent with its identification as an aggregation-prone region. Fibre diffraction and dye-binding analyses further show that the peptide adopts a homotypic steric-zipper interface closely resembling topological features observed in the chorion filament structures, where cA_8-13_-mediated zippers form lateral packing interfaces. In addition, preformed cA_8-13_ aggregates can template the assembly of chorion filaments. This interpretation is further supported by multimeric AF predictions that prioritize homotypic cA_8-13_ steric zippers as early intermediates and mirrors recent structural studies revealing analogous roles for APRs in pathological tau assemblies^66^. This involvement of short motifs in amyloid nucleation and stabilization is well established for pathological amyloids^74-76^, where they frequently form steric zippers that define protofilament interfaces^57^ and contribute to polymorphic behaviour^66,77^. Similarly, recent high-resolution structures of antimicrobial functional amyloids demonstrate that short amyloidogenic segments can encode self-complementary interfaces that stabilize the assembled state^25-28^. Our findings extend these concepts to the chorion biological system.

Beyond the cA_8-13_ motif, energetic analysis pinpoints two additional fibril-core stabilizing segments (residues 32-38 and 45-50) that form heterotypic steric-zipper interfaces within the central protofilament. Thermodynamic profiling of the resolved polymorphs revealed that in Type 3 filaments, stabilizing contributions remain concentrated within the discrete APRs, whereas in Type 1 and 5 polymorphs, stabilization becomes redistributed across the central protofilament core. Together with the extensive structural similarity between polymorphs and their coexistence within the same filament trajectories, these observations suggest that the resolved conformers occupy closely related states within a shared assembly landscape rather than distinct folds. In this framework, cA_8-13_-mediated interactions may provide permissive early assembly interfaces that bias formation toward the observed β-serpentine scaffold, while additional APR-derived contacts stabilize increasingly consolidated protofilament organizations. An alternative possibility is that some polymorphs correspond to partially consolidated structural states sampled during filament maturation under the assembly conditions. In such a scenario, the shared structural scaffold and progressive redistribution of stabilizing interactions across Type 3, Type 1, and Type 5 filaments would be consistent with increasingly consolidated protofilament architectures, in line with previous hierarchical assembly models proposed for amyloid fibril structures^78^ and maturation-associated structural remodeling described in other amyloids^65-67^.

Despite forming a well-defined cross-β architecture, chorion filaments exhibit relatively low thermodynamic stability and dissociate at modest denaturant concentrations. This behaviour is characteristic of many functional amyloids^48,57,79,80^, which often lack the deeply buried hydrophobic cores typical of pathological filaments and instead rely on distributed stabilization across multiple interfaces. Such an arrangement may enable the chorion matrix to combine mechanical robustness with the capacity for dynamic remodelling during oocyte development. Together, our findings support a model in which chorion amyloid assembly is governed by a modular and hierarchical mechanism in which APR-derived steric zippers nucleate assembly and promote secondary nucleation, while additional interfaces stabilize mature protofilament architectures. More broadly, these results demonstrate that assembly principles commonly associated with pathological amyloids, including secondary nucleation, APR-driven interfaces, and hierarchical stabilization, can also operate within biologically regulated functional amyloid systems.

## Materials and Methods

### Sequence Retrieval and Curation

Central domains of Lepidopteran chorion proteins were retrieved from LepChorionDB^81^ and UniProt^82^. Homologs were collected across 32 species spanning moths and butterflies. Sequences were aligned using Clustal Omega (v1.2.4) with default parameters and manually inspected in Jalview (v2.11.5.1) to ensure correct register of the central domain. The consensus sequence archetype (_1_SYGGEGIGNV_10-11_AVAGELPVAG_20_-_21_KTAVAGRVPI_30-31_IGAVGFGGPA_40-41_GAAGAVSIAGR_51_) was defined from previous work^38,43,52^ and the positional consensus of the alignment.

### Phylogenetic and Correlation Analysis

A maximum-likelihood phylogeny was reconstructed from the aligned sequences using IQ-TREE2^83^ (v2.3.2) under the LG+G4 model, selected by Bayesian information criterion. Branch support was assessed with 1000 ultrafast bootstrap replicates. The resulting tree was visualized in circular layout using the ggtree R package (v3.10.1), and tip annotations were added from species metadata. For each sequence, the fraction of glycine, proline, aromatic residues, and charged residues was calculated using in-house R scripts. Net charge at pH 7 and hydropathy (GRAVY index) were estimated using the Peptides package (v2.4.4). Correlations among all features were computed using corrplot (v0.92), with significance tested by pairwise Pearson correlation (p < 0.05).

### Sequence Dissimilarity and Dimensionality Reduction

Pairwise distances were computed as

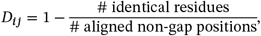

yielding an *n* × *n* distance matrix over the alignment. Classical multidimensional scaling (MDS) was applied (cmdscale, k = 50), followed by Uniform Manifold Approximation and Projection (UMAP, v0.2.10.0) to embed sequences in two dimensions. Resulting coordinates were plotted using ggplot2 (v3.5.1), with colour coding by species or sequence-derived parameters. Each aligned sequence was compared to the archetype using position-wise BLOSUM62 substitution scores. Average values were computed over all non-gap sites and used to generate the similarity gradient.

### Computational Predictions and Analysis

Fibril stability was quantified using our previously established thermodynamic profiling pipeline^4,84-86^, in which all inter-residue interaction energies within the cryo-EM-defined amyloid core were computed and summed to yield per-residue stabilization contributions. Aggregation-prone regions within chorion sequences were predicted using TANGO^87^ and CORDAX^88^. TANGO predicts β-aggregation propensity based on a simple statistical physicochemical principles of secondary structure formation extended by the assumption that the core regions of an aggregate are fully buried under defined physicochemical conditions^74^, whereas CORDAX identifies sequence regions compatible with cross-β fibril architectures through structural threading and energetic evaluation against experimentally determined amyloid folds^76^. AF3 predictions were performed using the AlphaFold Server^89^. Structural inspection, model manipulation, and molecular graphics were performed in ChimeraX^90^ (v1.10.1). All statistical analyses and data visualization were carried out in GraphPad Prism (v10.6.1).

### Peptide Synthesis

Peptides were synthesized on a MultiPep 2 robot (CEM) using Fmoc solid-phase synthesis with N-acetylated/C-amidated termini. Purity (>90%) was confirmed by HPLC (Shimadzu Nexera) and LC–MS (Shimadzu LCMS-2050) and peptides were ether-precipitated and stored at –20 °C (**Fig. S2**).

### Cryo-EM Grid Preparation

The cA peptide was dried under a nitrogen stream, resuspended in Milli-Q water at a concentration of 17.5 mg/mL, and incubated for 7 days to allow aggregation. After 7 days, and immediately before grid preparation, the peptide solution was diluted 1:1 in Milli-Q water. A suspension of the fibril sample (3.5 **µ**L) was applied to glow-discharged holey carbon grids (Quantifoil Cu R1.2/1.3, 300 mesh) and rapidly frozen in liquid ethane using an FEI Vitrobot Mark IV (Thermo Fischer Scientific). The grids were loaded onto a 300 kV Titan Krios microscope (Thermo Fischer Scientific), and movies were acquired on a Falcon 4i direct electron detector in EER mode, with a pixel size of 0.73485 Å and a dose of 1 e^−^/Å^2^ per frame. The defocus range was −0.5 to −2.5 µm. A total of 5,541 movies were collected.

### Helical Reconstruction

Micrograph movies were motion-corrected and dose-weighted using the motion-correction implementation in RELION^91,92^, and the contrast transfer function (CTF) was estimated from each motion-corrected micrograph using CTFFIND-4.1^93^. Micrographs were first filtered using figure-of-merit (FOM) values calculated by template-matching the characteristic 4.7 Å cross-β Fourier signal across each input micrograph^94^. Micrographs were then selected on the skewness of their FOM-image pixel-value distribution (rlnMicrographScoreSkewness ≥ 1.0), retaining 4,051 of 5,541 micrographs (73%) in which a high-FOM amyloid signal separated clearly from the Gaussian background; the remaining micrographs lacked detectable amyloid features. The retained micrographs were re-picked with filament tracing enabled (filament length 300 pixels, width 100 pixels, picking threshold 0.4, carbon-edge detection threshold 0.9), yielding 2,939,488 picked segments. Picked segments were initially extracted with a box size of 768 pixels and rescaled 6-fold to 128 pixels (effective pixel size 4.41 Å/px) for initial two-dimensional (2D) classification using a 540Å mask diameter. Successive rounds of 2D classification facilitated polymorph identification and separation where particles representing specific polymorphs were combined, re-extracted using a 384-box size without rescaling were subject to 3D refinement. A de novo 3D initial model was generated with the relion_helix_inimodel2d program^95^ from the selected 2D class averages. The initial model was used as the reference for a first 3D auto-refinement; a soft-edged solvent-flattened mask (low-pass filtered to 15 Å) was then generated from the output of this refinement and applied during a second 3D auto-refinement. Helical rise and twist were initially imposed and then optimised within a narrow search range, converging to rise/twist values of 4.784 Å / −1.800° (Type 1), 4.751 Å / −2.821° (Type 3), 4.861 Å / −1.645° (Type 4), 4.788 Å / −1.465° (Type 5) and 4.814 Å / −0.691° (Type 6). Bayesian polishing^96^ and three rounds of CTF refinement^97^ were then performed on the per-polymorph particle sets, followed by a final round of 3D auto-refinement with amyloid-specific Blush regularisation^94^. Masking and post-processing with relion_helix_toolbox^95^ were carried out with a central Z length of 30%, imposing C1 symmetry. The final maps reached resolutions of 2.66 Å, 3.03 Å, 3.00 Å, 3.07 Å, and 4.03 Å, respectively, using an FSC criterion of 0.143 (**Table S1** and **Fig. S4**).

### Model Building and Refinement

Atomic models for Type 1, 3, and 5 polymorphs were initially generated with ModelAngelo^97^, using the archetype sequence. A single helical rung was extracted from each ModelAngelo output in ChimeraX^90^ and expanded to a 5-layer model using the refined helical rise and twist; the 5-layer models were then subjected to real-space refinement against the experimental maps in ISOLDE^98^. Refinement and validation were performed in Phenix^99^. Type 4 and 6 polymorphs were not modelled because de novo model building was ambiguous. Validation statistics for each modelled polymorph are reported in Table S1.

### X-ray diffraction

The cA_8-13_ peptide was dried under a nitrogen stream and resuspended in Milli-Q water at 20mg/ml and left to aggregate for 7 days. A fibril suspension (6 µL) was deposited between two wax-coated capillaries positioned closely and mounted horizontally on a glass slide. X-ray diffraction data were collected from the aligned fibre using a Rigaku MicroMax-003 high-brilliance X-ray generator, equipped with a Rigaku HyPix direct photon detector, incident radiation Cu K-alpha. The specimen-to-film distance was set at 77 mm. Exposure time was set to 90 s. Still images were recorded using the program ChrysalisPro and subsequently displayed and measure using iMosflm (v7.4.0)^100^. The Clearer software was used to generate simulated diffraction patterns^101^.

### Fluorescence Aggregation Assays

Aliquots of the archetype were dried under a nitrogen stream, pretreated with 1,1,1,3,3,3-hexafluoro-2-propanol (HFIP; Sigma-Aldrich), dried again under nitrogen, and resuspended in Milli-Q water. First, cA kinetics were measured by resuspending films to a concentration of 110µM, followed by 1:1 serial dilution to the indicated concentrations. Non-linear fittings were performed using AmyloFit^102^. For seeding assays, the films were resuspended in PBS to a final concentration of 220 µM and mixed with 5% (v/v) preformed seeds generated by sonication for 12 min at 65% amplitude using alternating 30 s on/off cycles in a QSonica bath sonicator. Thioflavin-T (ThT) was added to a final concentration of 25 µM in 96-well plates (Greiner #675096). Fluorescence was measured at 37 °C in a CLARIOstar plate reader (BMG Labtech). Cross-seeding assays were fitted using prebuilt functions available in GraphPad Prism (v10.6.1).

### Thermodynamic Stability Measurements

Chemical depolymerization and thermodynamic stability of chorion filaments was determined using urea-induced disassembly coupled to flow-induced dispersion analysis (FIDA) of the soluble fraction (Fida1, FidaBio). Preformed sonicated filaments (100 µM monomer equivalent) were incubated for at least 16 h at 25 °C in Milli-Q containing increasing concentrations of urea (0-5.4 M). Following incubation, samples were analysed by FIDA to quantify the residual monomer fraction. The analysis was performed using the method below (**Table 1**) with a 100cm length and 75µm inner diameter capillary. For each elution profile, the fraction of soluble monomer was calculated as the ratio of the area under the curve (AUC) corresponding to the monomer peak divided by the total integrated AUC for that sample. This value reflects the proportion of protein that remains in the soluble state under each denaturant condition. The AUC for both monomer peak and total area was calculated with Fityk^103^ and an two-state depolymerization model^104^.

**Table 1.** FIDA method for analysis of amyloid fibril stability.

| Step | Component | Pressure | Time | Temp (°C) |
| --- | --- | --- | --- | --- |
| Wash | 1M NaOH | 3500 | 45 | 25 |
| Wash | Water | 3500 | 45 | 25 |
| Equilibration | Buffer | 3500 | 30 | 25 |
| Sample | Protein | 75 | 20 | 25 |
| Measurement | Buffer | 1500 | 120 | 25 |
| Wash | Water | 3500 | 45 | 25 |

### Transmission Electron Microscopy

Ten microliters of each sample were applied to glow-discharged copper grids (Formvar/Carbon 400-mesh; Electron Microscopy Sciences) for 3 min. Grids were washed with Milli-Q water and stained with 2% (w/v) uranyl acetate for 1 min. Samples were imaged using a JEOL 1400 Plus transmission electron microscope operated at 80 keV.

### Fluorescence Spectroscopy and Dimensionality Reduction

Fluorescence spectroscopy measurements were performed using preformed filaments diluted to 300 µM with PBS and a final pFTAA concentration of 0.5 µM. Measurements were done in a low-volume black 384-well plate (Corning) using a CLARIOStar plate reader (BMG Labtech). After excitation at 440 nm, the emission spectra were measured between 465 nm and 700 nm. Principal component analysis was performed using the PCA package available in R.

## Acknowledgements

We thank Sjors Scheres for constructive discussions and feedback on the manuscript, and Sofia Lövestam for feed-back on cryo-EM analysis. We dedicate this work to the memory of ***Stavros J. Hamodrakas***, whose pioneering structural and biophysical studies of silkmoth chorion proteins laid the conceptual foundations for understanding the molecular organization of insect egg coats, and to ***Fotis C. Kafatos***, whose seminal early contributions established the molecular and evolutionary framework of chorion biology that made this work possible.

NL and the Louros lab were supported by a scholarship from the Thomas O. Hicks Scholar in Medical Research. KK was partially supported by the O’Donnell Brain Institute (OBI) Sprouts grant program. Computational resources were provided by the BioHPC cluster supported by the Lyda Hill Department of Bioinformatics at UTSW. We also acknowledge the assistance of the UT Southwestern Electron Microscopy Core, funded by the NIH grants 1S10OD021685-01A1. We thank the Cryo-EM core facility (CEMF) at UT Southwestern Medical Center, for support with cryo-EM studies. We thank the Structural Biology Laboratory at UT Southwestern Medical Center for support with X-ray diffraction studies. Research reported in this publication was supported by the Office Of The Director, National Institutes of Health of the National Institutes of Health under Award Number S10OD025018. The content is solely the responsibility of the authors and does not necessarily represent the official views of the National Institutes of Health.

## Author contributions

NL conceived and initiated the study. KK, PK, and HT performed the experiments. NL performed computational analyses. KK, PK, and NL performed data analysis. MID, and NL provided resources and funding. NL supervised the project. KK, PK, and NL prepared the original draft, and all authors contributed to the review and editing of the final version of this manuscript.

## Data Availability statement

All experimental and computational raw data generated are available from the corresponding author on reasonable request.

