## Supplementary Information for "A functional amyloid scaffold shapes insect egg coats"

This PDF contains:

Supplementary Table 1

Supplementary Figures 1-4

**Supplementary Table 1.** Cryo-EM data collection, refinement, and validation statistics.

| **Data Collection** | Type 1 | Type 3 | Type 4 | Type 5 | Type 6 |
|---|---|---|---|---|---|
| Electron Microscope | Titan Krios | | | | |
| Nominal Magnification | 130,000x | | | | |
| Voltage (kV) | 300 | | | | |
| Detector | Falcon 4i | | | | |
| Dose per Frame (e-/A^2) | 1 | | | | |
| Defocus Range (μm) | -1 to -2.5 μm | | | | |
| Pixel Size (A) | 0.73485 | | | | |
| **Reconstruction** | | | | | |
| Total number of micrographs | 5,541 | | | | |
| Total particles extracted | 2,939,488 | | | | |
| Particles after 2D classification | 964,559 | 379,686 | 358,399 | 251,841 | 88,766 |
| Particles in final reconstruction | 370,481 | 213,614 | 124,369 | 94,152 | 34,543 |
| Map Resolution FSC=0.143 (Å) | 2.66 | 3.03 | 3 | 3.07 | 4.03 |
| Helical Rise (Å) | 4.784 | 4.751 | 4.861 | 4.788 | 4.814 |
| Helical Twist (degrees) | -1.8 | -2.8 | -1.6 | -1.5 | -0.7 |
| Symmetry Imposed | C1 | C1 | C1 | C1 | C1 |
| Map Sharpening B-factor (Å^2) | -75.5 | -88.6 | -60.8 | -70 | -60.2 |
| **Refinement** | | | | | |
| Initial Model | ModelAngelo | ModelAngelo | --- | ModelAngelo | --- |
| Refinement Software | ISOLDE/Phenix | ISOLDE/Phenix | --- | ISOLDE/Phenix | --- |
| Model-Map Resolution FSC=0.5 (Å) | 2.49 | 2.97 | --- | 3.1 | --- |
| **Model composition** | | | | | |
| Non-hydrogen Atoms | 1,785 | 1,135 | --- | 1,880 | --- |
| Protein Residues | 290 | 195 | --- | 310 | --- |
| Ligands | 0 | 0 | --- | 0 | --- |
| Mean B-factor Protein (Å^2) | 84.3 | 97.3 | --- | 73.33 | --- |
| **Validation** | | | | | |
| MolProbity Score | 1.96 | 1.8 | --- | 1.76 | --- |
| Clashscore | 4.84 | 12.89 | --- | 5.45 | --- |
| Poor Rotamers (%) | 3.57 | 0 | --- | 0 | --- |
| **Ramachandran plot** | | | | | |
| Favored (%) | 95.83 | 96.97 | --- | 92.59 | --- |
| Allowed (%) | 4.17 | 3.03 | --- | 7.41 | --- |
| Outliers (%) | 0 | 0 | --- | 0 | --- |
| **R.m.s deviations** | | | | | |
| RMS Bond Lengths (A) | 0.012 | 0.011 | --- | 0.012 | --- |
| RMS Bond Angles (deg) | 1.94 | 1.82 | --- | 1.81 | --- |

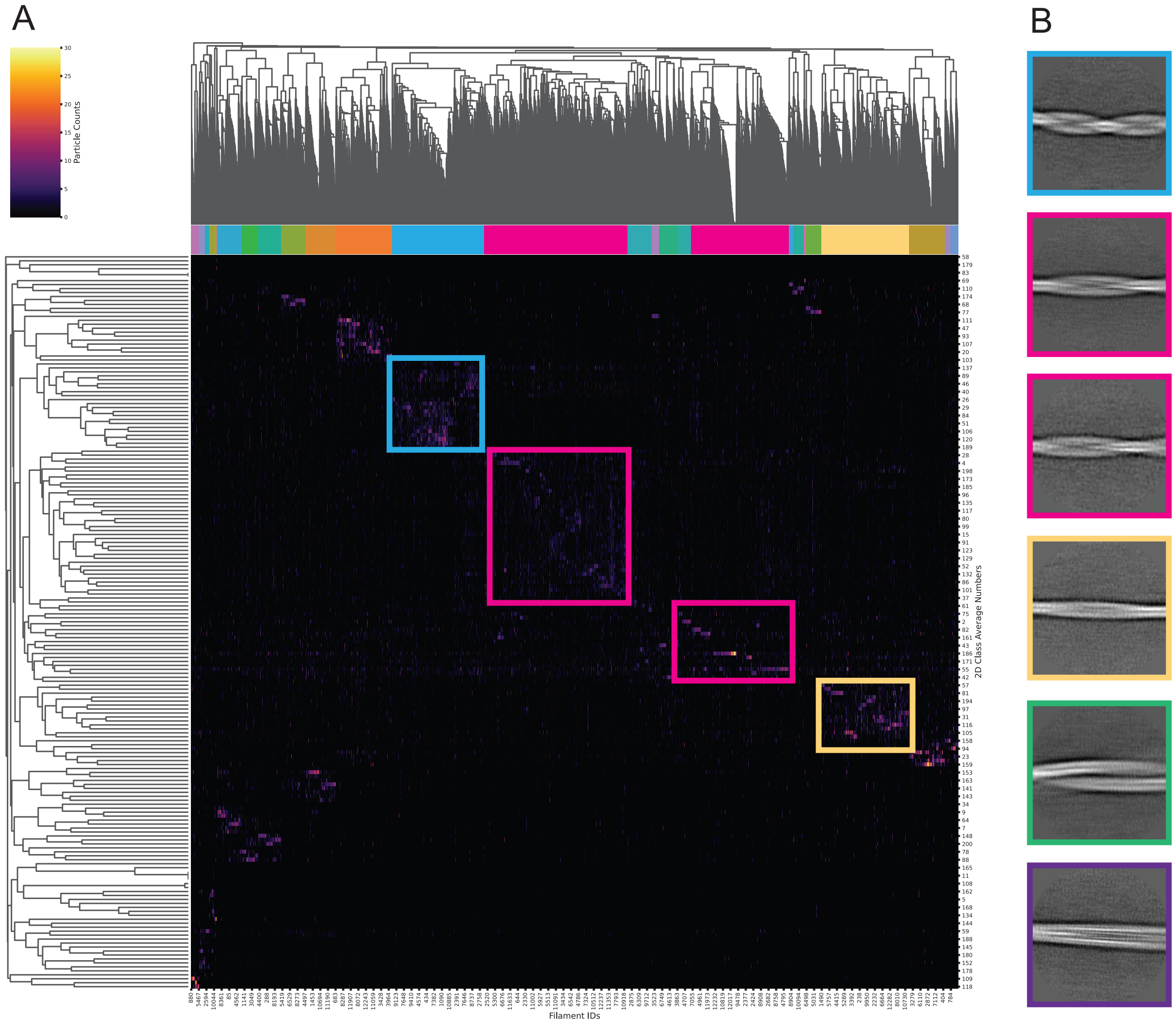


**Supplementary Figure 1. Hierarchical clustering of cA polymorphs. (A)** Hierarchical classification of individual filament segments according to their assigned 2D class average (vertical) and the picked filament ID (horizontal). **(B)** 2D class averages of filament types.

A

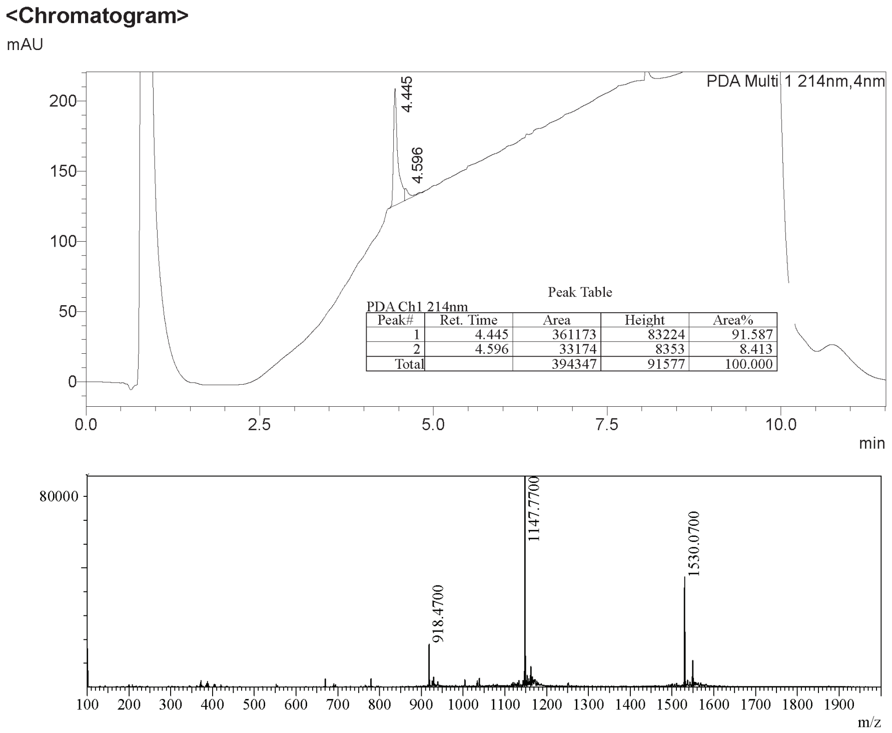


Peak Table

PDA Ch1 214nm

| Peak# | Ret. Time | Area | Height | Area% |
|---|---|---|---|---|
| 1 | 4.445 | 361173 | 83224 | 91.587 |
| 2 | 4.596 | 33174 | 8353 | 8.413 |
| Total | | 394347 | 91577 | 100.000 |

B

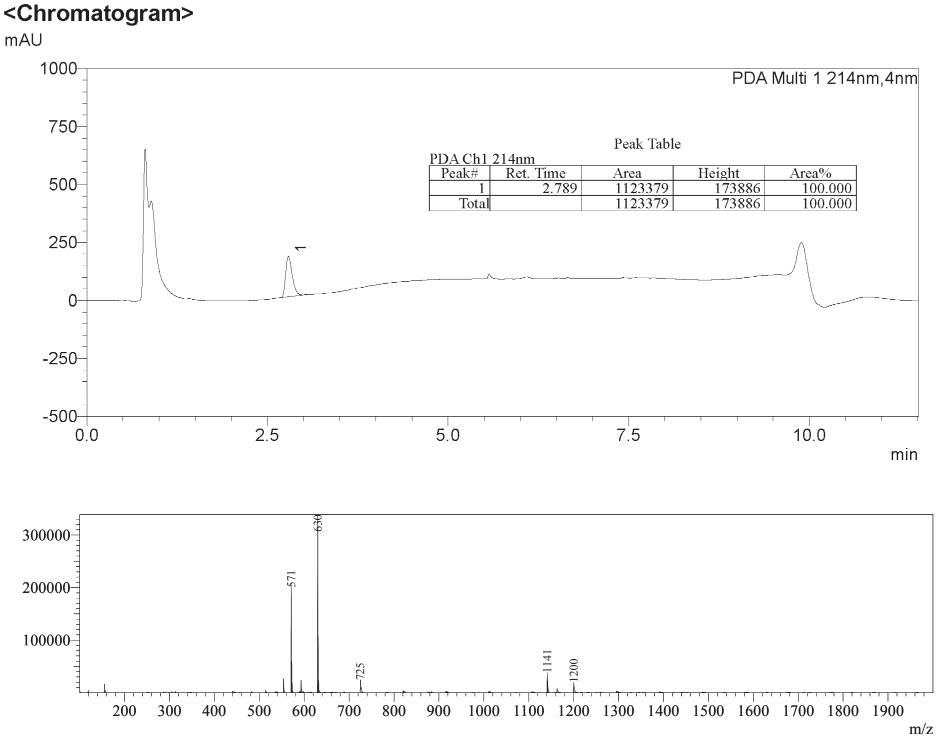


Peak Table

PDA Ch1 214nm

| Peak# | Ret. Time | Area | Height | Area% |
|---|---|---|---|---|
| 1 | 2.789 | 1123379 | 173886 | 100.000 |
| Total | | 1123379 | 173886 | 100.000 |

**Supplementary Figure 2. LC-MS characterization of the cA and $cA_{8-13}$ synthetic peptides. (A)** Representative LC-MS chromatogram and mass spectrometry analysis for cA. **(B)** Representative LC-MS chromatograms and mass spectrometry analysis for the $cA_{8-13}$ peptide.

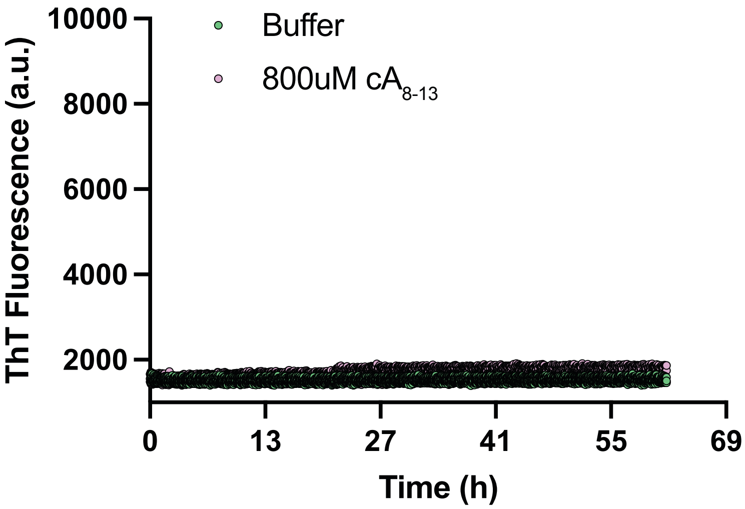


**Supplementary Figure 3. Fluorescence aggregation kinetics of the $cA_{8-13}$ motif.** No ThT signal was detected despite confirmation of amyloid fibril formation.

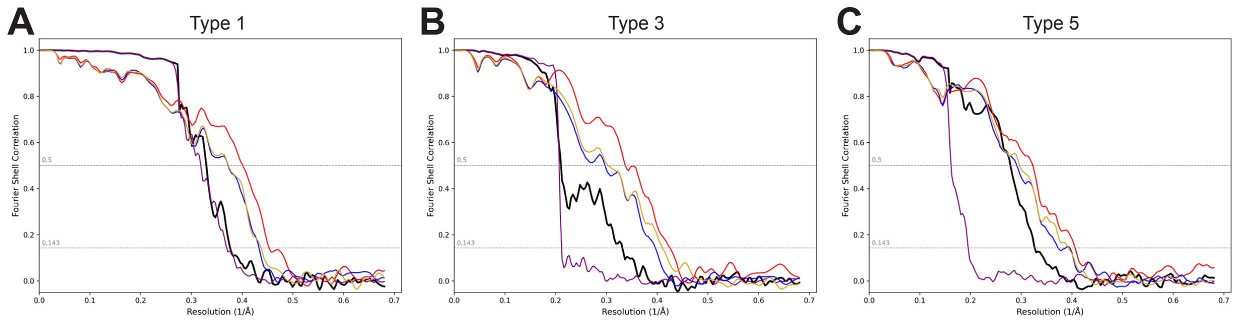


**Supplementary Figure 4. Resolution estimates for the chorion fibril datasets.** For each reconstructed polymorph, solvent-corrected Fourier-shell correlation (FSC) curves between independently refined half-maps are shown in black. FSC curves between the refined atomic model and against the post-processed map are shown in red. FSC curves between the model refined against half-map 1 are shown in blue, FSC curves of the same model against half-map 2 are shown in yellow, and phase randomized curves of the independently refined half-maps are shown in purple. Dotted lines indicate the FSC thresholds of 0.143 for half-map resolution estimates and 0.5 for the model-to-map estimates.
